# A Membrane-Bound Cytochrome Enables *Methanosarcina acetivorans* to Conserve Energy to Support Growth from Extracellular Electron Transfer

**DOI:** 10.1101/590380

**Authors:** Dawn E Holmes, Toshiyuki Ueki, Hai-Yan Tang, Jinjie Zhou, Jessica A Smith, Gina Chaput, Derek R Lovley

**Affiliations:** Department of Microbiology, University of Massachusetts Amherst, Morrill IV N Science Center, Amherst, MA 01003, USA; Department of Physical and Biological Sciences, Western New England University, Springfield, MA, 01119, USA; Jiangsu Provincial Key Lab for Organic Solid Waste Utilization, National Engineering Research Center for Organic-based Fertilizers, Jiangsu Collaborative Innovation Center for Solid Organic Waster Resource Utilization, Nanjing Agricultural University, Nanjing, 210095, China; School of Life Science and Biotechnology, Dalian University of Technology, Dalian, Liaoning Province, China, 116024; Department of Biology, American International College, Springfield, MA

**Keywords:** anaerobic respiration, extracellular electron transfer, anthraquinone-2,6,-disulfonate reduction, AQDS reduction, Rnf complex, *c*-type cytochrome, methanogen, archaea

## Abstract

Conservation of energy to support growth solely from extracellular electron transfer was demonstrated for the first time in a methanogen. *Methanosarcina acetivorans* grew with methanol as the sole electron donor and the extracellular electron acceptor anthraquione-2,6-disulfonate (AQDS) as the sole electron acceptor when methane production was inhibited with bromoethanesulfonate. Transcriptomics revealed that transcripts for the gene for the transmembrane, multi-heme, *c*-type cytochrome MmcA were 4-fold higher in AQDS-respiring cells versus methanogenic cells. A strain in which the gene for MmcA was deleted failed to grow via AQDS reduction whereas strains in which other cytochrome genes were deleted grew as well as the wild-type strain. The MmcA-deficient strain grew with the conversion of methanol or acetate to methane, suggesting that MmcA has a specialized role as a conduit for extracellular electron transfer. Enhanced expression of genes for methanol conversion to methyl-coenzyme M and components of the Rnf complex suggested that methanol is oxidized to carbon dioxide in AQDS-respiring cells through a pathway that is similar to methyl-coenezyme M oxidation in methanogenic cells. However, during AQDS respiration the Rnf complex and reduced methanophenazine probably transfer electrons to MmcA, which functions as the terminal reductase for AQDS reduction. Extracellular electron transfer may enable survival of methanogens in dynamic environments in which oxidized humic substances and Fe(III) oxides are intermittently available. The availability of tools for genetic manipulation of *M. acetivorans* makes it an excellent model microbe for evaluating *c*-type cytochrome-dependent extracellular electron transfer in Archaea.

**Importance:** Extracellular electron exchange in *Methanosarcina* species and closely related Archaea plays an important role in the global carbon cycle and can enhance the speed and stability of anaerobic digestion, an important bioenergy strategy. The potential importance of *c*-type cytochromes for extracellular electron transfer to syntrophic bacterial partners and/or Fe(III) minerals in some Archaea has been suspected for some time, but the studies with *Methanosarcina acetivorans* reported here provide the first genetic evidence supporting this hypothesis. The results suggest parallels with Gram-negative bacteria, such as *Shewanella* and *Geobacter* species, in which outer-surface *c*-type cytochromes are an essential component for electrical communication with the extracellular environment. *M. acetivorans* offers an unprecedented opportunity to study mechanisms for energy conservation from the anaerobic oxidation of one-carbon organic compounds coupled to extracellular electron transfer in Archaea with implications not only for methanogens, but possibly also for anaerobic methane oxidation.

## Introduction

Extracellular electron exchange is central to the environmental function of diverse Archaea that oxidize and/or produce methane. Some methane-producing microorganisms can divert electron transfer from methane production to the reduction of extracellular electron carriers such as Fe(III), U(VI), V(IV), and anthraquinone-2,6-disulfonate (AQDS), a humic acid analog (1–9). Diversion of electron flux from methane production to extracellular electron transfer may influence the extent of methane production and metal geochemistry in anaerobic soils and sediments. Methanogens such as *Methanothrix* (formerly *Methanosaeta*) and *Methanosarcina* species can accept electrons via direct interspecies electron transfer from electron-donating partners, such as *Geobacter* species in important methanogenic environments such as anaerobic digesters and rice paddy soils (10–12). Anaerobic methane oxidation also plays an important role in the global carbon cycle and diverse anaerobic methane-oxidizing archaea (ANME) transfer electrons derived from methane oxidation to extracellular electron acceptors, such as other microbial species, Fe(III), or extracellular quinones (13–19). The electrical contacts for extracellular electron exchange have yet to be definitively identified in any of these Archaea.

It has been hypothesized that outer-surface cytochromes enable electron transfer to electron-accepting microbial partners or Fe(III) in some ANME (13–19). Genes for multi-heme *c*-type cytochromes that are present in ANME genomes can be highly expressed and in some instances the proteins have been detected. The putative function of outer-surface cytochromes is terminal electron transfer to extracellular electron acceptors, similar to the role that outer surface *c*-type cytochromes play in extracellular electron transfer in Gram-negative bacteria such as *Shewanella* and *Geobacter* species (20–22). Similar *c*-type cytochrome electrical contacts have been proposed for Fe(III)-reducing Archaea such as *Ferroglobus* and *Geoglobus* species (23, 24). However, the study of the mechanisms for extracellular electron transfer in these archaea has been stymied by the lack of microorganisms available in pure culture that can grow via extracellular electron transfer and are genetically tractable.

Tools are available for genetic manipulation of the methanogen *Methanosarcina acetivorans* (25–27). A methyl-coenzyme M reductase from an uncultured ANME was introduced into *M. acetivorans* to generate a strain that could convert methane to acetate with simultaneous reduction of Fe(III) (28). Most of the electrons from the methane consumed were recovered in acetate (28) and it was not shown that energy was conserved from Fe(III) reduction. *In vitro* reactions catalyzed by membrane vesicles of wild-type *M. acetivorans* suggested that the membrane-bound heterodisulfide reductase HdrDE reduced Fe(III)-citrate and AQDS, and that an outer-surface multi-heme *c*-type cytochrome, might also function as a potential electron donor for Fe(III)-citrate reduction (29). However, *in vitro* assays with cell components are not a definitive approach for determining the physiologically relevant mechanisms involved in the reduction of Fe(III) and AQDS because many reduced co-factors and redox-proteins, including *c*-type cytochromes, can non-specifically reduce these electron acceptors (30). Analysis of the phenotypes of intact cells that result from specific gene deletions can provide more conclusive evidence.

Here we report that *M. acetivorans* can be grown in the absence of methane production with AQDS as the sole electron acceptor. Analysis of gene expression patterns and phenotypes of gene deletion strains suggest a mechanism for energy conservation during extracellular electron transfer.

## Results and Discussion

### Growth of *M. acetivorans* with AQDS as the sole terminal electron acceptor

In medium with methanol provided as the electron donor and AQDS as a potential electron acceptor, *M. acetivorans* simultaneously produced methane and reduced AQDS (Figure 1a). The addition of bromoethanesulfonate (BES) inhibited methane production and increased the extent of AQDS reduction (Figure 1b; Supplementary Figure S1). Metabolism of methanol (Figure 1c) was accompanied by cell growth (Figure 1d). In the BES-amended cultures 6.2 mM methanol was consumed with the reduction of 15.7 mM AQDS. When the need to divert some of the methanol metabolized to cell biomass is considered, this stoichiometry is consistent with the oxidation of methanol to carbon dioxide with AQDS serving as the sole electron acceptor: CH_3_OH + 3AQDS + H_2_O → 3AH_2_QDS + CO_2_.

**Figure 1.**
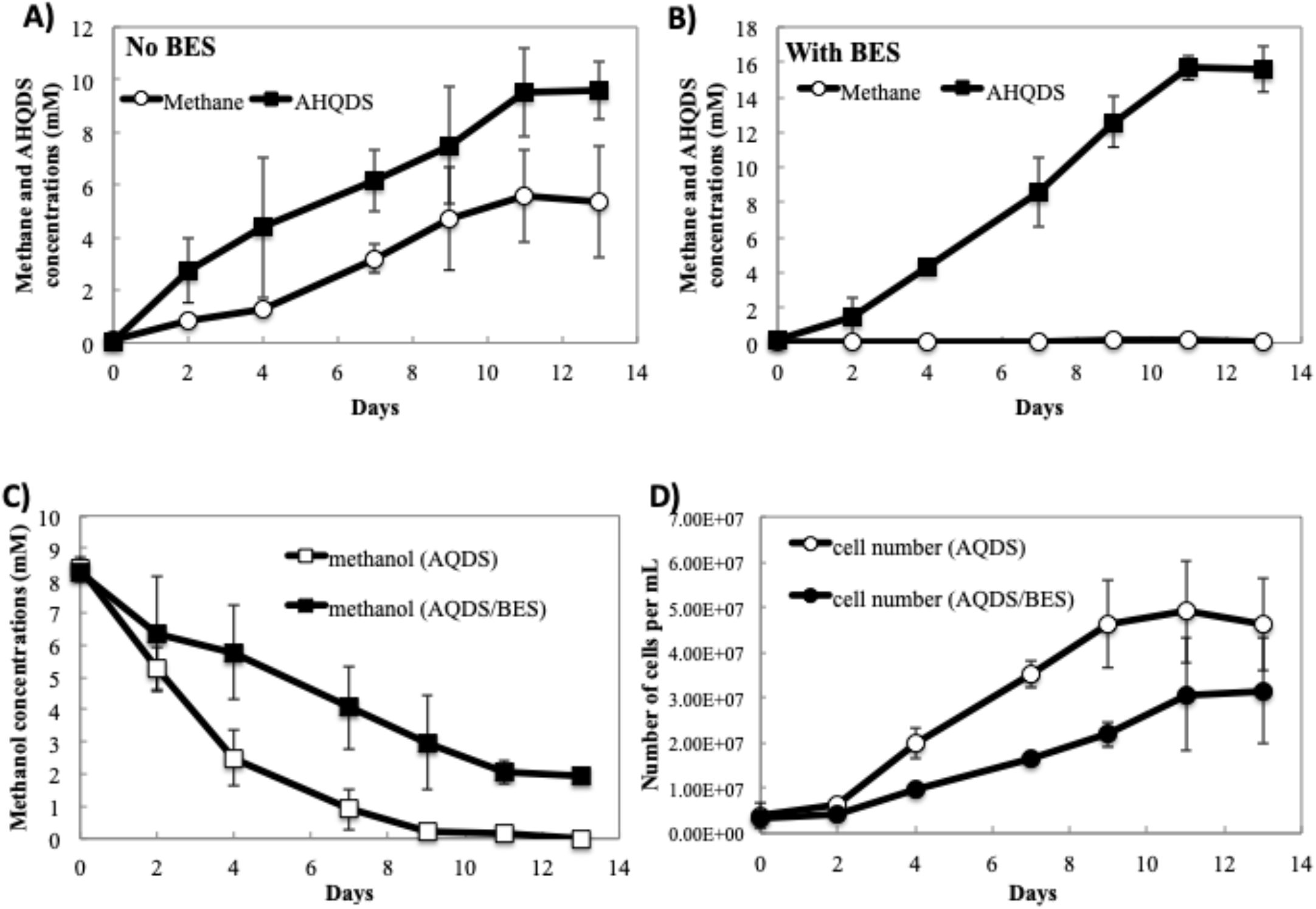
Growth of *Methanosarcina acetivorans* with methanol provided as an electron donor and AQDS as an electron acceptor in the presence or absence of BES. (A) Methane and AHQDS concentrations generated by cultures grown without BES; (B) Methane and AHQDS concentrations generated by cultures grown with BES; (C) Methanol concentrations and (D) cell numbers from cultures grown in the presence or absence of BES. The complete inhibition of methane production in the presence of BES is also shown on an expanded scale in Supplementary Figure S1.

The greater consumption of methanol in the absence of BES (Figure 1c), was in accordance with the extent of AQDS reduction and the simultaneous conversion of methanol to methane: 4CH_3_OH → 3CH_4_ + CO_2_ + 2H_2_O.

The methanol oxidation coupled with AQDS reduction in the presence of BES described here is the first demonstration of a methanogen conserving energy to support growth with electron transfer to an external electron acceptor as the sole means of energy conservation. The ability of *M. acetivorans* to grow in this manner, and the availability of tools for genetic manipulation (25–27) provide the opportunity for functional analysis of extracellular electron transfer in an archaeon.

### Transcriptomics and gene deletion studies demonstrate that the multi-heme *c*-type cytochrome MmcA is important for AQDS reduction

In order to obtain insight into potential electron carriers involved in AQDS reduction, the transcriptome of cells grown with AQDS as the sole electron acceptor in the presence of BES was compared with the transcriptome of cells grown with methanol in the absence of AQDS or BES, so that methane production was the sole route of electron flux. The median log_2_ RPKM value for the cells grown via methanogenesis (5.2) was substantially higher than for the cells grown via AQDS reduction (4.0). These results are consistent with the finding that cells grown via methanogenesis grew ∼4 times faster than cells respiring AQDS (generation time for AQDS-respiring cells was 3 days vs 0.7 days for methanogenic cells).

Remarkably, despite the overall lower transcription rate of cells grown via AQDS reduction, the transcripts for gene MA0658, which encodes a seven-heme, outer-surface *c*-type cytochrome, were 4-fold higher in AQDS-reducing versus methanogenic cells (Table 1, Supplementary Table S1A). For future reference, this cytochrome was designated MmcA (<u>m</u>embrane <u>m</u>ulti-heme <u>c</u>ytochrome A). Multi-heme *c*-type cytochromes are of particular interest as potential electron carriers in extracellular electron transport because of the well-documented role of multi-heme *c*-type cytochromes in bacteria such as *Shewanella* and *Geobacter* species that are highly effective in extracellular electron transfer (20–22). MA3739, a gene coding for a five-heme *c*-type cytochrome, was transcribed at similar levels as *mmcA*, and 4 fold more transcripts were detected in AQDS-reducing than methanogenic cells (Table 1).

**Table 1.** Differential expression of genes coding for *c*-type cytochrome proteins in *M. acetivorans* cells grown with methanol provided as the electron donor and AQDS as the electron acceptor in the presence of BES, or cells grown via methanogenesis with methanol as the substrate. Genes were only considered differentially expressed if the fold change was ≥ 2 and the P-value and FDR (False Discovery Rate) were <0.05. NS: no significant difference in read abundance between conditions

| Locus ID | # heme groups | # transmembrane helices | Predicted Localization | Fold up-regulated in AQDS/BES vs methanogenesis | P-value | FDR |
| --- | --- | --- | --- | --- | --- | --- |
| MA0658 | 7 | 1 | Membrane | 3.95 | 0.002 | 0.006 |
| MA3739 | 5 | 0 | Unknown | 4.14 | 0.009 | 0.02 |
| MA0167 | 1 | 1 | Membrane | 5.97 | 0.002 | 0.006 |
| MA2925 | 2 | 1 | Membrane | 1.21 (NS) | 0.29 | 0.37 |
| MA2908 | 2 | 1 | Membrane | 1.03 (NS) | 0.87 | 0.89 |

There are three other putative *c*-type cytochrome genes in the *M. acetivorans* genome (31). Transcripts for MA0167, which encodes a mono-heme cytochrome with predicted localization in the cell membrane, were 6-fold more abundant in cells grown via AQDS respiration (Table 1). Functional analysis of the outer-membrane of *G. sulfurreducens* has suggested that a mono-heme *c*-type cytochrome may play a role in regulating the expression of multi-heme *c*-type cytochromes, possibly by providing a sensor function (32, 33). It is possible that the protein encoded by MA0167 is playing a similar role in *M. acetivorans*. The number of transcripts for MA2925 and MA2908, both of which encode two-heme *c*-type cytochromes, was comparable in AQDS-reducing versus methanogenic cells (Table 1). These cytochromes are homologous to methylamine utilization protein G (MauG) and di-heme cytochrome c peroxidase (CcpA). MauG is required for aerobic methylamine metabolism (34–36), and CcpA proteins reduce hydrogen peroxide to water and protect the cell from reactive oxygen species (37, 38). Thus, it seems unlikely that either of these cytochromes is involved in extracellular electron transfer.

In order to evaluate the potential role of *c*-type cytochromes in AQDS reduction, deletion mutant strains were constructed in *M. acetivorans* for each *c*-type cytochrome gene in the genome (Table 1). Only the deletion of *mmcA* inhibited AQDS reduction (Figure 2a). Deletion of *mmcA* had a slight impact on methanogenic growth with methanol (Figure 2b).

**Figure 2.**
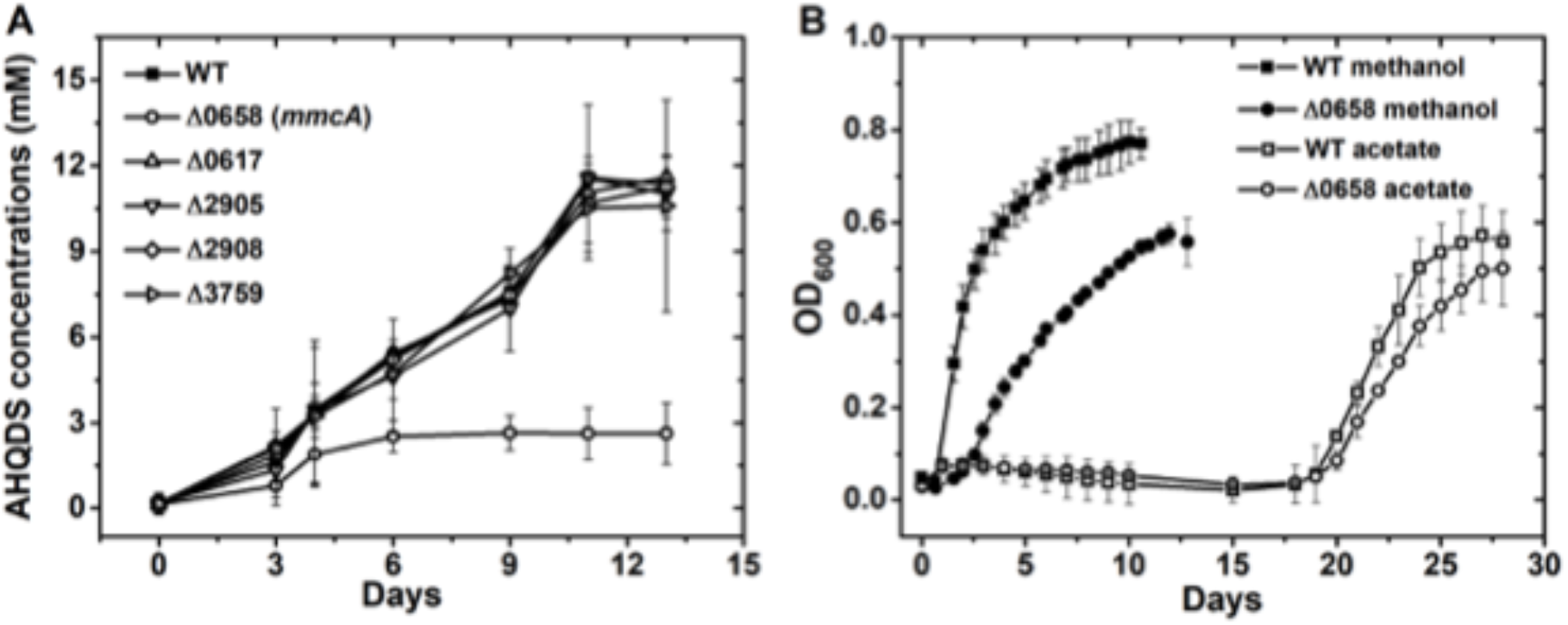
Impact of deletion of *c*-type cytochrome genes on growth of *M. acetivorans* under different conditons. (A) AHQDS production during growth with methanol as the electron donor and AQDS as the acceptor in the presence of BES. The locus ID for the deleted cytochrome gene is designated next to the corresponding symbol. (B) Growth of wild-type and ΔMA0658 strains under methanogenic conditions as measured by A_600_ with methanol or acetate provided as substrates.

These results suggest that MmcA is an essential component for extracellular electron transfer to AQDS, but not for the conversion of methanol to methane. This conclusion was further supported by the finding that *mmcA* was highly transcribed in AQDS-reducing cells, however, its expression levels were below the median log_2_ RPKM values for methanogenic cells (Table 1 and Supplementary Table S1).

Previous studies have suggested that MmcA is part of the Rnf complex, which is required for acetoclastic methanogenesis (39) and that it is co-transcribed with Rnf genes located in the same region of the chromosome (40). However, deletion of the MmcA gene did not substantially impact growth on acetate (Figure 2B) or transcription of other genes from the Rnf complex (Supplementary Figure S2). Furthermore, the expression profiles of *mmcA* and genes for the Rnf complex were also different (Tables 1 and 2).

**Table 2.** Comparison of transcripts from genes coding for components of the Rnf and Mrp complexes in *M. acetivorans* cells grown with methanol and AQDS in the presence of BES, or cells grown via methanogenesis with methanol. egative values indicate that genes were more significantly expressed in methanogenic cells. Genes were only considered differentially expressed if the fold change was ≥ 2 and the P-value and FDR (False Discovery Rate) were <0.05. NS: no significant difference in read abundance

| Locus ID | Annotation | Gene | Fold up-regulated in AQDS/BES vs methanogenesis | P-value | FDR |
| --- | --- | --- | --- | --- | --- |
| MA0659 | electron transport complex protein RnfC | *rnfC* | 1.52 (NS) | 0.02 | 0.04 |
| MA0660 | electron transport complex protein RnfD | *rnfD* | 1.23 (NS) | 0.19 | 0.26 |
| MA0661 | electron transport complex protein RnfG | *rnfG* | 1.66 (NS) | 0.006 | 0.01 |
| MA0662 | electron transport complex protein RnfE | *rnfE* | 1.45 (NS) | 0.02 | 0.05 |
| MA0663 | electron transport complex protein RnfA | *rnfA* | 1.66 (NS) | 0.006 | 0.01 |
| MA0664 | electron transport complex protein RnfB | *rnfB* | 1.57 (NS) | 0.008 | 0.01 |
| MA4572 | multisubunit sodium/proton antiporter, MrpA subunit | *mrpA* | 5.44 | $8.87 \times 10^{-8}$ | $8.13 \times 10^{-6}$ |
| MA4665 | multisubunit sodium/proton antiporter, MrpB subunit | *mrpB* | 5.41 | $1.57 \times 10^{-7}$ | $1.07 \times 10^{-5}$ |
| MA4570 | multisubunit sodium/proton antiporter, MrpC subunit | *mrpC* | 6.50 | $1.21 \times 10^{-7}$ | $9.14 \times 10^{-6}$ |
| MA4569 | multisubunit sodium/proton antiporter, MrpD subunit | *mrpD* | 4.84 | $2.05 \times 10^{-7}$ | $1.18 \times 10^{-5}$ |
| MA4568 | multisubunit sodium/proton antiporter, MrpE subunit | *mrpE* | 3.70 | $6.32 \times 10^{-6}$ | $8.86 \times 10^{-5}$ |
| MA4567 | multisubunit sodium/proton antiporter, MrpF subunit | *mrpF* | 4.79 | $5.21 \times 10^{-7}$ | $1.86 \times 10^{-5}$ |
| MA4566 | multisubunit sodium/proton antiporter, MrpG subunit | *mrpG* | 4.57 | $5.70 \times 10^{-7}$ | $1.98 \times 10^{-5}$ |

### Model for Electron Transport to AQDS via MmcA

MmcA is a strong candidate for the terminal AQDS reductase because its localization in the cell membrane (40) is likely to provide access to AQDS and because of the well-known role of outer-membrane multi-heme *c*-type cytochromes in reduction of AQDS and various forms of Fe(III) in Gram-negative bacteria such as *Shewanella* and *Geobacter* species (20–22, 41). It was previously suggested that MmcA could be a terminal reductase for the reduction of soluble Fe(III)-citrate, based on the *in vitro* oxidation of MmcA in membrane vesicles upon addition of Fe(III)-citrate (29). Such *in vitro* assays can be poor predictors of *in vivo* activity because Fe(III)-citrate typically oxidizes *c*-type cytochromes *in vitro*, regardless of physiological function, due to its very positive redox potential. However, as detailed below, multiple lines of evidence support a model in which energy can be conserved when MmcA serves as the terminal reductase during methanol oxidation coupled to AQDS reduction (Figure 3).

**Figure 3.**
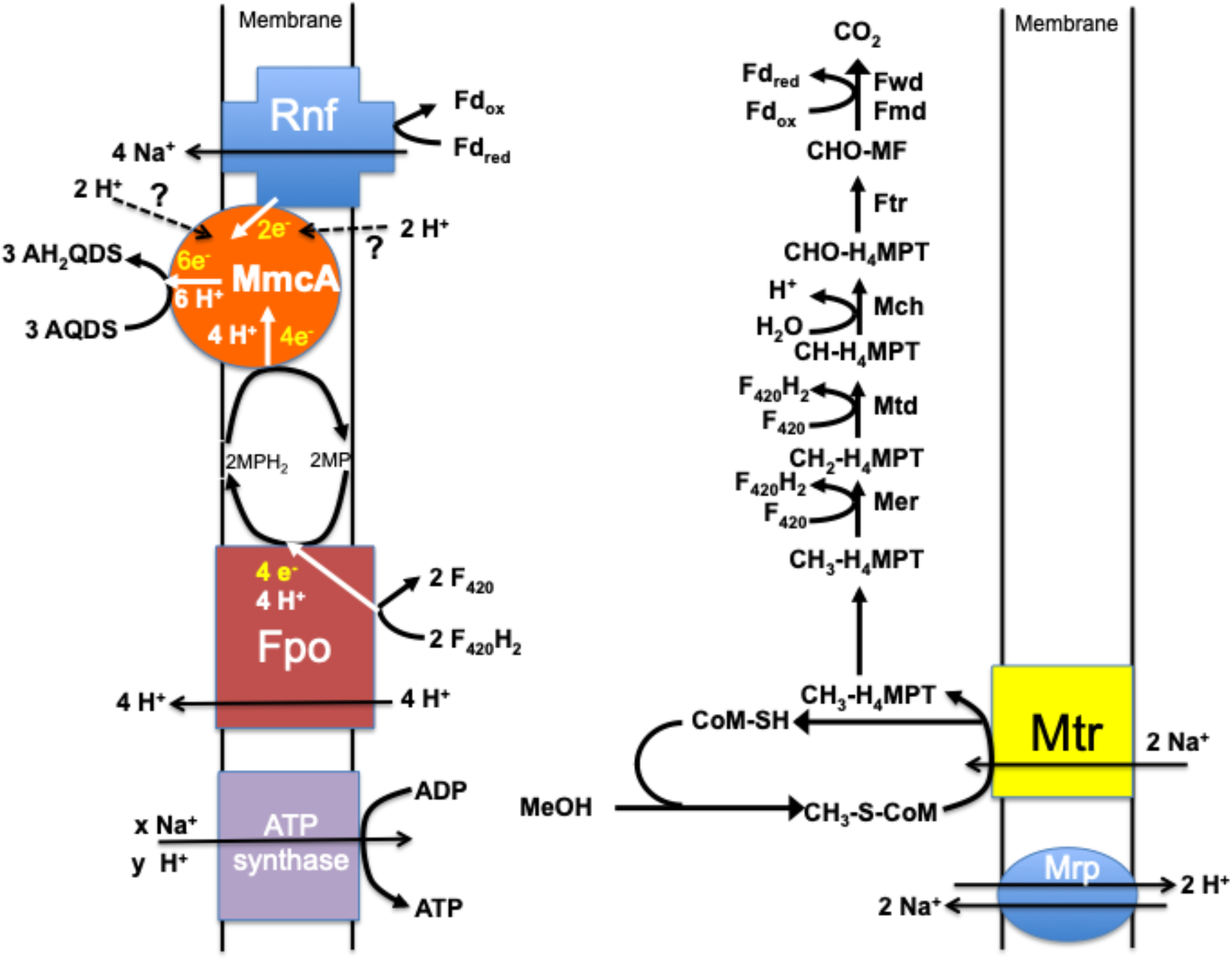
Proposed model for extracellular electron transport to AQDS by *Methanosarcina acetivorans* when methanol is provided as the electron donor and methanogenesis is prevented with the addition of BES.

During methane production from methanol, methanol is converted to CH_3_-CoM by the activity of three enzymes, methyltransferase 1 (MtaB), methyltransferase 2 (MtaA), and methanol corrinoid protein (MtaC) (42–44). The oxidation of one molecule of CH_3_-CoM to CO_2_ generates the reducing equivalents necessary to reduce three molecules of CH_3_-CoM to methane. During methanol oxidation coupled to AQDS reduction in the presence of BES, the step that reduces CH_3_-CoM to methane is blocked, but the option for CH_3_-CoM oxidation remains (Figure 3). Genes coding for enzymes involved in the oxidation of CH_3_-CoM to carbon dioxide were more highly expressed in methanogenic cells, consistent with increased overall transcriptional activity in methanogenic cells and the need for this pathway to generate reductants to support methanogenesis (Supplementary Table S2). However, transcription of genes coding for enzymes involved in CH_3_-CoM oxidation were also well above the median log_2_ RPKM value in the AQDS-respiring cells, suggesting that this pathway is also important for methanol oxidation coupled to AQDS reduction (Supplementary Tables S1A).

Differential expression of genes encoding isomers of MtaB, MtaA, and MtcC suggested that there might be some differences in the route for methanol conversion to CH_3_-CoM (Table 3). The genes for the isomers MtaB1, MtaA1, and MtaC1 were more highly expressed in methanogenic cells, whereas AQDS-respiring cells had higher transcript abundance for genes coding for the alternative MtaB, MtaA, and MtaC isomers (Table 3). Differences in the activity of these isomers are unknown, but in previous studies *mtaA1, mtaB1*, and *mtaC1* genes were specifically transcribed during methanogenesis from methanol and MtaA1 was required for growth on methanol, whereas MtaA2 was dispensable (44).

**Table 3.** Differential expression of genes coding for methanol methyltransferase enzymes in *M. acetivorans* cells grown with methanol provided as an electron donor and AQDS provided as an electron acceptor in the presence of BES or cells grown via methanogenesis with methanol. Negative values indicate that genes were more significantly expressed in methanogenic cells. Genes were only considered differentially expressed if the fold change was ≥ 2 and the P-value and FDR (False Discovery Rate) were <0.05. NS: no significant difference in read abundance

| Locus ID | Annotation | Gene | Fold up-regulated in AQDS/BES vs methanogenesis | P-value | FDR |
| --- | --- | --- | --- | --- | --- |
| MA4379 | Co-methyl-5-hydroxybenzimidazolylcobamide:2-mercapto-ethanesulphonic acid methyltransferase, isozyme 1 | *mtaA1* | -1.68 | 0.008 | 0.02 |
| MA0455 | methanol:5-hydroxybenzimidazolylcobamide methyltransferase, isozyme 1 | *mtaB1* | -6.84 | 0.002 | 0.007 |
| MA0456 | corrinoid-containing methyl-accepting protein, isozyme 1 | *mtaC1* | -7.95 | 0.001 | 0.005 |
| MA4392 | methanol:5-hydroxybenzimidazolylcobamide methyltransferase, isozyme 2 | *mtaB2* | 68.55 | $1.53 \times 10^{-10}$ | $6.88 \times 10^{-7}$ |
| MA4391 | corrinoid-containing methyl-accepting protein, isozyme 2 | *mtaC2* | 48.28 | $8.34 \times 10^{-10}$ | $1.25 \times 10^{-6}$ |
| MA1615 | Co-methyl-5-hydroxybenzimidazolylcobamide:2-mercapto-ethanesulphonic acid methyltransferase, isozyme 2 | *mtaA2* | 5.39 | $3.40 \times 10^{-7}$ | $1.50 \times 10^{-5}$ |
| MA1616 | methanol:5-hydroxybenzimidazolylcobamide methyltransferase, isozyme 3 | *mtaB3* | 9.66 | $7.45 \times 10^{-8}$ | $7.56 \times 10^{-6}$ |
| MA1617 | corrinoid-containing methyl-accepting protein, isozyme 3 | *mtaC3* | 8.49 | $3.78 \times 10^{-7}$ | $1.55 \times 10^{-5}$ |

Oxidation of methanol to carbon dioxide is expected to yield reduced ferredoxin and reduced F_420_ (F_420_H_2_). It is likely that the Rnf complex oxidizes reduced ferredoxin with electron transfer to MmcA (45). Despite the lower overall gene transcript abundance in AQDS-respiring cells, transcripts for genes coding for components of the Rnf complex were slightly higher than those in methanogenic cells (Table 2), suggesting an important role for the Rnf complex in energy conservation from methanol oxidation coupled to AQDS reduction.

In methanogenic cells the membrane-bound Fpo complex (F_420_:methanophenazine oxidoreductase) oxidizes F420H2 derived from methanol oxidation with the reduction of methanophenazine and proton extrusion (46–50). Transcript abundance for all Fpo subunit genes was higher in methanogenic cells than AQDS-reducing cells, as expected because of the importance of Fpo in oxidizing F_420_H_2_ in cells producing methane and the overall higher gene expression levels in methanogenic cells (Supplementary Table S3). However, the number of transcripts for all of the Fpo complex genes was significantly higher than the median log_2_ RPKM value in AQDS-respiring cells (Supplementary Table S1A), suggesting that Fpo is important for the oxidation of F_420_H_2_ generated in methanol-oxidizing, AQDS-reducing cells. The reduced methanophenazine that Fpo generates from F_420_H_2_ oxidation can transfer electrons to MmcA (39, 40, 45, 51). Although it has also been proposed that reduced methanophenazine may be able to directly transfer electrons to extracellular electron carriers in *M. acetivorans* (29), the requirement for MmcA for growth via AQDS reduction indicates that this is an unlikely route for AQDS reduction.

In methanogenic cells, reduced methanophenazine can also donate electrons to the membrane-bound heterodisulfide reductase HdrDE (50, 52–55). *In vitro* evidence with membrane-vesicles suggested that HdrDE can reduce AQDS with CoM-SH and CoB-SH oxidation to form CoM-S-S-CoB (29). However, the redox-active components of HdrDE responsible for electron transfer to an electron acceptor are localized to the cytoplasmic side of the membrane (50) and thus unlikely to access extracellular AQDS *in vivo*. The relative expression of *hdrD* and *hdrE* was slightly lower in AQDS-reducing cells than methanogenic cells (Supplementary Table S1). Furthermore, the inability of the MmcA-deficient strain to grow via AQDS reduction indicates that HdrDE is not capable of functioning as the sole AQDS reductase to support growth. Thus, in the lack of strong evidence for a role for HdrDE, the likely simpler and more direct route for AQDS-dependent oxidation of reduced methanophenazine is electron transfer to MmcA.

From these considerations, and the current understanding of the function of the redox proteins involved (50, 56, 57), a positive balance of Na^+^ and H^+^ outside the cell to support the generation of ATP during AQDS respiration is possible (Figure 3). In this model two Na^+^ must be translocated into the cell for the initial oxidation of CH_3_-S-CoM. Two moles of F_420_H_2_ and one mole of reduced ferrodoxin are generated per mole of CH3-S-CoM oxidized to carbon dioxide. Fpo oxidizes the F_420_H_2_ with H^+^ extrusion and the reduction of methanophenazine. The reduced methanophenazine transfers electrons to MmcA, which reduces AQDS. The Rnf complex oxidizes the reduced ferredoxin coupled with Na^+^ translocation and the reduction of MmcA. MmcA may transfer protons as well as electrons during AQDS reduction as observed in other *c*-type cytochromes (58–63). The ATP synthase couples both Na^+^ and H^+^ transport to ATP synthesis (64), but the H^+^/Na^+^ antiporter complex Mrp can be important for balancing external Na^+^/H^+^ ratios (65). Genes for Mrp were highly expressed in AQDS-reducing cells (Table 2).

Uncertainties in the stoichiometry of Na^+^/H^+^ transport per ATP synthesized and the total amount of H^+^ translocated prevent an accurate estimate of the theoretical ATP yield per mole of methanol oxidized with the reduction of AQDS. However, it is clear that net ATP synthesis is likely from the proposed metabolic route, consistent with the observed growth of *M. acetivorans* with methanol oxidation coupled to AQDS reduction.

### Implications

The discovery that *M. acetivorans* can conserve energy to support growth from the oxidation of a one-carbon compound coupled to the reduction of an extracellular electron acceptor has important implications for the biogeochemistry of anaerobic soils and sediments and provides a genetically tractable model microbe for further analysis of the mechanisms of extracellular electron transfer in *Archaea*. Humic substances and Fe(III) are often abundant extracellular electron acceptors in a wide variety of anaerobic soils and sediments and their availability for microbial respiration can reduce the extent of methane production (66–69). Competition for electron donors between methanogens and Fe(III)- and humics-reducing microorganisms is one factor (70, 71). However, the finding that some methanogens may conserve energy by reducing extracellular electron acceptors suggests a mechanism for methanogens to survive in environments in which Fe(III) and oxidized forms of humic substances are abundant and then rapidly switch to methane production as these extracellular electron acceptors are depleted.

A comprehensive survey of the ability of diverse methanogens to conserve energy to support growth from electron transport to extracellular electron acceptors is warranted. Most methanogens, including other *Methanosarcina* species, lack membrane-bound multi-heme cytochromes like MmcA and would need other mechanisms for extracellular electron transfer. The finding that MmcA is not essential for methane production, and that expression of *mmcA* was increased when AQDS served as an electron acceptor, suggests that the primary role of MmcA is extracellular electron transfer. If so, the presence of MmcA in *M. acetivorans* further suggests that there are environments in which the capacity for extracellular electron transfer substantially benefits *M. acetivorans*.

A wide diversity of archaea are capable of extracellular electron transfer (72), but the mechanisms are poorly understood. For archaea such as *Ferroglobus placidus* (23), *Geoglobus ahangari* (24), and diverse ANME (13–19) it has been proposed that outer-membrane cytochromes are the terminal reductase. The rapid non-physiological reduction of extracellular electron acceptors by a range of redox-active proteins and co-factors *in vitro* necessitates genetically tractable model organisms for physiologically relevant functional studies. Thus, *M. acetivorans* may serve as an important model organism for better understanding cytochrome-based extracellular electron transfer in Archaea.

## Materials and Methods

### Strains and growth conditions

*Methanosarcina acetivorans* strains were routinely cultured under strict anaerobic conditions at 37°C in the previously described (25) medium with either 8.5 mM methanol or 40 mM acetate provided as substrates.

*M. acetivorans* mutant strains were constructed with *M. acetivorans* WWM1 (Δ*hpt*) (73) as the parent strain as described previously (26). For construction of MA0658, MA3739, MA2908, MA0167, and MA2925 deletion strains, genes were replaced with the *pac* gene (puromycin resistance gene). First, regions 500-1000 bp upstream and downstream from the target genes were amplified by PCR (Supplementary Table S4). The DNA fragments of the upstream and downstream regions of MA0658 were digested with SacI/XbaI and EcoRI/XhoI. Upstream and downstream regions of MA3739 were digested with SalI/XbaI and SacI/NotI. Upstream and downstream regions of MA2908, MA0167, and MA2925 were digested with XhoI/HindIII and BamHI/NotI. The upstream fragment was ligated into the pJK3 plasmid (25). The downstream fragment was ligated into the pJK3 plasmid already containing the upstream fragment. This recombinant plasmid was then linearized and used for transformation. The deletion and replacement of all genes with *pac* was verified with primers (Supplementary Table S4). All transformants were selected on medium supplemented with puromycin (2 μM final concentration), as previously described (25).

Additions of anthraquinone-2,6,-disulphonate (AQDS) were made from a concentrated stock to provide a final concentration of 16 mM. Cysteine was omitted from all cultures. When noted, 2-bromoethanesulfonate (BES) was added from a concentrated stock to provide a final concentration of 15 mM. Growth with AQDS was measured by determining numbers of cells stained with acridine orange with epifluorescence microscopy (74). For comparing methanogenic growth in wild-type and mutant cells, growth was monitored by spectrometry at an absorbance of 600 nm (75).

### Analytical techniques

Methanol concentrations were monitored with a gas chromatograph equipped with a headspace sampler and a flame ionization detector (Clarus 600; PerkinElmer Inc., CA). Methane in the headspace was measured by gas chromatography with a flame ionization detector (Shimadzu, GC-8A) as previously described (76). Production of reduced AQDS reduction was monitored by spectrophotometry at an absorbance of 450 nm as previously described (77).

### RNA extraction

Cells were harvested from triplicate 50 mL cultures of *M. acetivorans* grown with methanol (10 mM) provided as the electron donor and AQDS (16 mM) in the presence of the methanogenesis inhibitor BES (15 mM) or via methanogenesis with 40 mM methanol provided as substrate. Cells were split into 50 mL conical tubes (BD Sciences), mixed with RNA Protect (Qiagen) in a 1:1 ratio, and pelleted by centrifugation at 3,000 x g for 15 minutes at 4°C. Pellets were then immediately frozen in liquid nitrogen and stored at −80°C. Total RNA was extracted from all six cell pellets according to the previously described protocol (78) and cleaned with the RNeasy Mini Kit (Qiagen). All RNA samples were then treated with Turbo DNA-free DNase (Ambion, Austin, TX). In order to ensure that samples were not contaminated with genomic DNA, PCR with primers targeting the 16S rRNA gene was done with RNA that had not been reverse transcribed. Further enrichment of mRNA was done with the MICROB*Express* kit (Ambion), according to the manufacturer’s instructions.

### RT-PCR analysis

Total RNA was prepared from *M. acetivorans* hpt and ΔMA0658 strains grown methanogenically with acetate (40 mM). Complementary DNA (cDNA) was prepared by reverse transcription with AMV reverse transcriptase (New England Biolabs, MA) with primers TCAGCATGCCTCATTCCAAC (MA0659) or TCGCAGACAGCCTTAACGTC (MA0664) according to the manufacturers specifications. This cDNA was then used as a template for PCR with the following primer pairs: CAGTGACCTCGCTTATGTCC/TCAGCATGCCTCATTCCAAC (MA0695) or TGTGGAGGTTGCGGATTTGC/TCGCAGACAGCCTTAACGTC (MA0664). The amplified fragments were analyzed by agarose gel electrophoresis.

### Illumina sequencing and data analysis

Directional multiplex libraries were prepared with the ScriptSeq™ v2 RNA-Seq Library Preparation Kit (Epicentre) and paired end sequencing was performed on a Hi-Seq 2000 platform at the Deep Sequencing Core Facility at the University of Massachusetts Medical School in Worchester, Massachusetts.

All raw data generated by Illumina sequencing were quality checked by visualization of base quality scores and nucleotide distributions with FASTQC (http://www.bioinformatics.babraham.ac.uk/proiects/fastqc/). Initial raw non-filtered forward and reverse sequencing libraries contained an average of 124,551,285 +/− 8,421,388 reads that were ∼100 basepairs long. Sequences from all of the libraries were trimmed and filtered with Trimmomatic (79) with the sliding window approach set to trim bases with quality scores lower than 3, strings of 3+N’s, and reads with a mean quality score lower than 20. Bases were also cut from the start and end of reads that fell below a threshold quality of 3, and any reads smaller than 50 bp were eliminated from the library. These parameters yielded an average of 115,861,910 +/− 2,278,492 quality reads per RNAseq library.

All paired-end reads were then merged with FLASH (80), resulting in 45,331,795 +/− 3,260,585 reads with an average read length of 145 basepairs. After merging the QC-filtered reads, SortMeRNA (81) was used to separate all ribosomal RNA (rRNA) reads from non-ribosomal reads.

### Mapping of mRNA reads

Trimmed and filtered mRNA reads from the triplicate samples for the two different culture conditions were mapped against the *M. acetivorans* strain C2A genome (NC_003552) downloaded from IMG/MER (img.jgi.doe.gov). Mapped reads were normalized with the RPKM (reads assigned per kilobase of target per million mapped reads) method (82, 83) using ArrayStar software (DNAStar). Analysis of reads from all three biological replicates for each condition demonstrated that results were highly reproducible. Therefore, all reported values were obtained after merging and averaging replicates. Expression levels were considered significant only when the log_2_ RPKM value was higher than that of the median log_2_ RPKM. Out of the 4721 predicted protein-coding genes in the *M. acetivorans* C2A genome, 2360 and 2362 had expression levels that were higher than the median in AQDS-respiring or methanogenic cells, respectively (Supplementary Table S1).

Reads were also normalized and processed for differential expression studies using the edgeR package in Bioconductor (84). Genes with p-values ≤ 0.05 and fold changes ≥ 2 were considered differentially expressed. Using these criteria, 827 genes were up-regulated and 778 genes were down-regulated in AQDS-respiring cells compared to methanogenic cells (Supplementary Table S5).

### Genome data analysis

Gene sequence data for *M. acetivorans* C2A was acquired from the US Department of Energy Joint Genome Institute (http://www.igi.doe.gov) or from Genbank at the National Center for Biotechnology Information (NCBI) (http://www.ncbi.nlm.nih.gov). Initial analyses were done with tools available on the Integrated Microbial Genomes (IMG) website (img.jgi.doe.gov). Some protein domains were identified with NCBI conserved domain search (85) and Pfam search (86) functions. Transmembrane helices were predicted with TMpred (87), TMHMM (88), and HMMTOP (89) and signal peptides were identified with PSORTb v. 3.0.2 (90) and Signal P v. 4.1 (91).

## Accession numbers

Illumina sequence reads have been submitted to the NCBI database under BioProject PRJNA501858 and submission number SUB4712594.

## Acknowledgments

This research was supported by the Army Research Office and was accomplished under Grant Number W911NF-17-1-0345. The views and conclusions contained in this document are those of the authors and should not be interpreted as representing the official policies, either expressed or implied, of the Army Research Office or the U.S. Government.

## References

1. Vargas M, Kashefi K, Blunt-Harris EL, Lovley DR. 1998. Microbiological evidence for Fe(III) reduction on early Earth. Nature 395:65–67.

2. Bond DR, Lovley DR. 2002. Reduction of Fe(III) oxide by methanogens in the presence and absence of extracellular quinones. Environ Microbiol 4:115–124.

3. Cervantes FJ, de Bok FAM, Tuan DD, Stams AJM, Lettinga G, Field JA. 2002. Reduction of humic substances by halorespiring, sulphate-reducing and methanogenic microorganisms. Environ Microbiol 4:51–57.

4. Bodegom PM, Scholten JC, Stams AJ. 2004. Direct inhibition of methanogenesis by ferric iron. FEMS Microbiol Ecol 49:261–8.

5. Liu D, Dong HL, Bishop ME, Wang HM, Agrawal A, Tritschler S, Eberl DD, Xie SC. 2011. Reduction of structural Fe(III) in nontronite by methanogen *Methanosarcina barkeri*. Geochim Cosmochim Acta 75:1057–1071.

6. Zhang J, Dong HL, Liu D, Fischer TB, Wang S, Huang LQ. 2012. Microbial reduction of Fe(III) in illite-smectite minerals by methanogen *Methanosarcina mazei*. Chem Geol 292:35–44.

7. Zhang J, Dong HL, Zhao LD, McCarrick R, Agrawal A. 2014. Microbial reduction and precipitation of vanadium by mesophilic and thermophilic methanogens. Chem Geol 370:29–39.

8. Sivan O, Shusta SS, Valentine DL. 2016. Methanogens rapidly transition from methane production to iron reduction. Geobiol 14:190–203.

9. Holmes DE, Orelana R, Giloteaux L, Wang LY, Shrestha P, Williams K, Lovley DR, Rotaru AE. 2018. Potential for methanosarcina to contribute to uranium reduction during acetate-promoted groundwater bioremediation. Microb Ecol 76:660–667.

10. Rotaru AE, Shrestha PM, Liu F, Markovaite B, Chen S, Nevin KP, Lovley DR. 2014. Direct interspecies electron transfer between *Geobacter metallireducens* and *Methanosarcina barkeri*. Appl Environ Microbiol 80:4599–605.

11. Rotaru AE, Shrestha PM, Liu FH, Shrestha M, Shrestha D, Embree M, Zengler K, Wardman C, Nevin KP, Lovley DR. 2014. A new model for electron flow during anaerobic digestion: direct interspecies electron transfer to *Methanosaeta* for the reduction of carbon dioxide to methane. Energy Environ Sci 7:408–415.

12. Holmes DE, Shrestha PM, Walker DJF, Dang Y, Nevin KP, Woodard TL, Lovley DR. 2017. Metatranscriptomic Evidence for Direct Interspecies Electron Transfer between *Geobacter* and *Methanothrix* Species in Methanogenic Rice Paddy Soils. Appl EnvironMicrobiol 83.

13. Myerdierks A, Kube M, Kostadinov I, Teeling H, Glockner FO, Reinhardt R, Amann R. 2010. Metagenome and mRNA expression analyses of anaerobic methanotrophic archaea of the ANME-1 group. Environ Microbiol 12:422–439.

14. McGlynn SE, Chadwick GL, Kempes CP, Orphan VJ. 2015. Single cell activity reveals direct electron transfer in methanotrophic consortia. Nature 526:531–U146.

15. Wegener G, Krukenberg V, Riedel D, Tegetmeyer HE, Boetius A. 2015. Intercellular wiring enables electron transfer between methanotrophic archaea and bacteria. Nature 526:587–590.

16. McGlynn SE. 2017. Energy metabolism during anaerobic methane oxidation in ANME Archaea. Microbes Environ 32:5–13.

17. Timmers PHA, Welte CU, Koehorst JJ, Plugge CM, Jetten MSM, Stams AJM. 2017. Reverse methanogenesis and respiration in methanotrophic archaea. Archaea 2017:1654237.

18. Cai C, Leu AO, Xie G-J, Guo J, Feng Y, Zhao J-X, Tyson GW, Yuan Z, Hu S. 2018. A methanotrophic archaeon couples anaerobic oxidation of methane to Fe(III) reduction. ISME J 12:1929–1939.

19. Krukenberg V, Riedel D, Gruber-Vodicka HR, Buttigieg PL, Tegetmeyer HE, Boetius A, Wegener G. 2018. Gene expression and ultrastructure of meso-and thermophilic methanotrophic consortia. Environ Microbiol 20:1651–1666.

20. Shi L, Dong H, Reguera G, Beyenal H, Lu A, Liu J, Yu H-Q, Fredrickson JK. 2016. Extracellular electron transfer mechanisms between microorganisms and minerals. Nat Rev Microbiol 14:651–662.

21. Ueki T, DiDonato LN, Lovley DR. 2017. Toward establishing minimum requirements for extracellular electron transfer in *Geobacter sulfurreducens*. FEMS Microbiol Lett 364:fnx093.

22. Aklujkar M, Coppi MV, Leang C, Kim BC, Chavan MA, Perpetua LA, Giloteaux L, Liu A, Holmes DE. 2013. Proteins involved in electron transfer to Fe(III) and Mn(IV) oxides by *Geobacter sulfurreducens* and *Geobacter uraniireducens*. Microbiol 159:515–35.

23. Smith JA, Aklujkar M, Risso C, Leang C, Giloteaux L, Holmes DE. 2015. Mechanisms involved in Fe(III) respiration by the hyperthermophilic archaeon *Ferroglobus placidus*. Appl Environ Microbiol 81:2735–44.

24. Manzella MP, Holmes DE, Rocheleau JM, Chung A, Reguera G, Kashefi K. 2015. The complete genome sequence and emendation of the hyperthermophilic, obligate iron-reducing archaeon “*Geoglobus ahangari*” strain 234(T). Stand Genomic Sci 10:77.

25. Metcalf WW, Zhang JK, Apolinario E, Sowers KR, Wolfe RS. 1997. A genetic system for Archaea of the genus *Methanosarcina:* liposome-mediated transformation and construction of shuttle vectors. Proc Natl Acad Sci U S A 94:2626–31.

26. Buan N, Kulkarni G, Metcalf W. 2011. Genetic methods for methanosarcina species. Methods Enzymol 494:23–42.

27. Nayak DD, Metcalf WW. 2017. Cas9-mediated genome editing in the methanogenic archaeon *Methanosarcina acetivorans*. Proc Natl Acad Sci U S A 114:2976–2981.

28. Soo VWC, McAnulty MJ, Tripathi A, Zhu F, Zhang L, Hatzakis E, Smith PB, Agrawal S, Nazem-Bokaee H, Gopalakrishnan S, Salis HM, Ferry JG, Maranas CD, Patterson AD, Wood TK. 2016. Reversing methanogenesis to capture methane for liquid biofuel precursors. Microb Cell Fact 15:11.

29. Yan Z, Joshi P, Gorski CA, Ferry JG. 2018. A biochemical framework for anaerobic oxidation of methane driven by Fe(III)-dependent respiration. Nature Comm 9:1642.

30. Coppi MV, O’Neil RA, Leang C, Kaufmann F, Methé BA, Nevin KP, Woodard TL, Liu A, Lovley DR. 2007. Involvement of *Geobacter sulfurreducens* SfrAB in acetate metabolism rather than intracellular Fe(III) reduction. Microbiol 153:3572–3585.

31. Kletzin A, Heimerl T, Flechsler J, van Niftrik L, Rachel R, Klingl A. 2015. Cytochromes c in Archaea: distribution, maturation, cell architecture, and the special case of *Ignicoccus hospitalis*. Front Microbiol 6.

32. Kim B-C, Leang C, Ding YR, Glaven RH, Coppi MV, Lovley DR. 2005. OmcF, a putative *c*-Type monoheme outer membrane cytochrome required for the expression of other outer membrane cytochrome in *Geobacter sulfurreducens*. J Bacteriol 187:4505–13.

33. Kim B-C, Postier BL, DiDonato RJ, Chaudhuri SK, Nevin KP, Lovley DR. 2008. Insights into genes involved in electricity generation in *Geobacter sulfurreducens* via whole genome microarray analysis of the OmcF-deficient mutant. Bioelectrochem 73:70–75.

34. Li X, Jones LH, Pearson AR, Wilmot CM, Davidson VL. 2006. Mechanistic possibilities in MauG-dependent tryptophan tryptophylquinone biosynthesis. Biochem 45:13276–83.

35. Pearson AR, Jones LH, Higgins L, Ashcroft AE, Wilmot CM, Davidson VL. 2003. Understanding quinone cofactor biogenesis in methylamine dehydrogenase through novel cofactor generation. Biochem 42:3224–30.

36. Wang Y, Graichen ME, Liu A, Pearson AR, Wilmot CM, Davidson VL. 2003. MauG, a novel diheme protein required for tryptophan tryptophylquinone biogenesis. Biochem 42:7318–25.

37. Hoffmann M, Seidel J, Einsle O. 2009. CcpA from *Geobacter sulfurreducens* is a basic di-heme cytochrome c peroxidase. J Mol Biol 393:951–65.

38. Atack JM, Kelly DJ. 2007. Structure, mechanism and physiological roles of bacterial cytochrome c peroxidases. Adv Microb Physiol 52:73–106.

39. Schlegel K, Welte C, Deppenmeier U, Muller V. 2012. Electron transport during aceticlastic methanogenesis by *Methanosarcina acetivorans* involves a sodium-translocating Rnf complex. FEBS J 279:4444–52.

40. Li Q, Li L, Rejtar T, Lessner DJ, Karger BL, Ferry JG. 2006. Electron transport in the pathway of acetate conversion to methane in the marine archaeon *Methanosarcina acetivorans*. J Bacteriol 188:702–10.

41. Voordeckers JW, Kim BC, Izallalen M, Lovley DR. 2010. Role of *Geobacter sulfurreducens* outer surface c-type cytochromes in reduction of soil humic acid and anthraquinone-2,6-disulfonate. Appl Environ Microbio 76:2371–2375.

42. Bose A, Pritchett MA, Rother M, Metcalf WW. 2006. Differential regulation of the three methanol methyltransferase isozymes in *Methanosarcina acetivorans* C2A. J Bacteriol 188:7274–83.

43. Galagan JE, Nusbaum C, Roy A, Endrizzi MG, Macdonald P, FitzHugh W, Calvo S, Engels R, Smirnov S, Atnoor D, Brown A, Allen N, Naylor J, Stange-Thomann N, DeArellano K, Johnson R, Linton L, McEwan P, McKernan K, Talamas J, Tirrell A, Ye W, Zimmer A, Barber RD, Cann I, Graham DE, Grahame DA, Guss AM, Hedderich R, Ingram-Smith C, Kuettner HC, Krzycki JA, Leigh JA, Li W, Liu J, Mukhopadhyay B, Reeve JN, Smith K, Springer TA, Umayam LA, White O, White RH, Conway de Macario E, Ferry JG, Jarrell KF, Jing H, Macario AJ, Paulsen I, Pritchett M, Sowers KR, et al. 2002. The genome of M. acetivorans reveals extensive metabolic and physiological diversity. Genome Res 12:532–42.

44. Bose A, Pritchett MA, Metcalf WW. 2008. Genetic analysis of the methanol-and methylamine-specific methyltransferase 2 genes of *Methanosarcina acetivorans* C2A. J Bacteriol 190:4017–26.

45. Wang M, Tomb JF, Ferry JG. 2011. Electron transport in acetate-grown *Methanosarcina acetivorans*. BMC Microbiol 11:165.

46. Kulkarni G, Kridelbaugh DM, Guss AM, Metcalf WW. 2009. Hydrogen is a preferred intermediate in the energy-conserving electron transport chain of *Methanosarcina barkeri*. PNAS 106:15915–15920.

47. Abken HJ, Deppenmeier U. 1997. Purification and properties of an F420H2 dehydrogenase from *Methanosarcina mazei* Go1. FEMS Microbiol Lett 154:231–237.

48. Baumer S, Ide T, Jacobi C, Johann A, Gottschalk G, Deppenmeier U. 2000. The F420H2 dehydrogenase from *Methanosarcina mazei* is a redox-driven proton pump closely related to NADH dehydrogenases. J Biol Chem 275:17968–17973.

49. Welte C, Deppenmeier U. 2011. Re-evaluation of the function of the F-420 dehydrogenase in electron transport of *Methanosarcina mazei*. FEBS J 278:1277–1287.

50. Welte C, Deppenmeier U. 2014. Bioenergetics and anaerobic respiratory chains of aceticlastic methanogens. Biochim Biophys Acta-Bioenergetics 1837:1130–1147.

51. Suharti S, Wang M, de Vries S, Ferry JG. 2014. Characterization of the RnfB and RnfG subunits of the Rnf complex from the archaeon *Methanosarcina acetivorans*. PLoS One 9:e97966.

52. Ide T, Baumer S, Deppenmeier U. 1999. Energy conservation by the H2:heterodisulfide oxidoreductase from *Methanosarcina mazei* Go1: identification of two proton-translocating segments. J Bacteriol 181:4076–80.

53. Heiden S, Hedderich R, Setzke E, Thauer RK. 1993. Purification of a cytochrome b containing H2:heterodisulfide oxidoreductase complex from membranes of Methanosarcina barkeri. Eur J Biochem 213:529–35.

54. Heiden S, Hedderich R, Setzke E, Thauer RK. 1994. Purification of a 2 subunit cytochrome-b containing heterodisulfide reductase from methanol grown *Methanosarcina barkeri*. Eur J Biochem 221:855–861.

55. Hedderich R, Hamann N, Bennati M. 2005. Heterodisulfide reductase from methanogenic archaea: a new catalytic role for an iron-sulfur cluster. Biol Chem 386:961–70.

56. Kumar VS, Ferry JG, Maranas CD. 2011. Metabolic reconstruction of the archaeon methanogen *Methanosarcina acetivorans*. BMC Syst Biol 5:28.

57. Benedict MN, Gonnerman MC, Metcalf WW, Price ND. 2012. Genome-scale metabolic reconstruction and hypothesis testing in the methanogenic archaeon *Methanosarcina acetivorans* C2A. J Bacteriol 194:855–65.

58. Yoshikawa S, Shimada A. 2015. Reaction mechanism of cytochrome c oxidase. Chem Rev 115:1936–89.

59. Morgado L, Dantas JM, Bruix M, Londer YY, Salgueiro CA. 2012. Fine tuning of redox networks on multiheme cytochromes from *Geobacter sulfurreducens* drives physiological electron/proton energy transduction. Bioinorg Chem Appl 2012:298739.

60. Louro RO, Catarino T, Turner DL, Picarra-Pereira MA, Pacheco I, LeGall J, Xavier AV. 1998. Functional and mechanistic studies of cytochrome c3 from *Desulfovibrio gigas:* thermodynamics of a “‘ proton thruster “‘. Biochem 37:15808–15.

61. Morgado L, Bruix M, Pessanha M, Londer YY, Salgueiro CA. 2010. Thermodynamic characterization of a triheme cytochrome family from *Geobacter sulfurreducens* reveals mechanistic and functional diversity. Biophys J 99:293–301.

62. Gunner MR, Mao J, Song Y, Kim J. 2006. Factors influencing the energetics of electron and proton transfers in proteins. What can be learned from calculations. Biochim Biophys Acta 1757:942–68.

63. Dantas JM, Morgado L, Aklujkar M, Bruix M, Londer YY, Schiffer M, Pokkuluri PR, Salgueiro CA. 2015. Rational engineering of *Geobacter sulfurreducens* electron transfer components: a foundation for building improved *Geobacter-based* bioelectrochemical technologies. Front Microbiol 6:752.

64. Schlegel K, Leone V, Faraldo-Gomez JD, Muller V. 2012. Promiscuous archaeal ATP synthase concurrently coupled to Na+ and H+ translocation. PNAS 109:947–952.

65. Jasso-Chavez R, Diaz-Perez C, Rodriguez-Zavala JS, Ferry JG. 2017. Functional Role of MrpA in the MrpABCDEFG Na+/H+ Antiporter Complex from the Archaeon *Methanosarcina acetivorans*. J Bacteriol 199.

66. Klupfel L, Piepenbrock A, Kappler A, Sander M. 2014. Humic substances as fully regenerable electron acceptors in recurrently anoxic environments. Nat Geosci 7:195–200.

67. Thamdrup B. 2000. Bacterial manganese and iron reduction in aquatic sediments. Adv Microb Ecol Vol 16 16:41–84.

68. Lovley DR. 1991. Dissimilatory Fe(III) and Mn(IV) reduction. Microbiol Rev 55:259–87.

69. Keller J, Weisenhorn P, Megonigal J. 2009. Humic acids as electron acceptors in wetland decomposition. Soil Biol Biochem 41:1518–1522.

70. Roden EE, Wetzel RG. 2003. Competition between Fe(III)-reducing and methanogenic bacteria for acetate in iron-rich freshwater sediments. Microb Ecol 45:252–258.

71. Lovley DR, Phillips EJP. 1987. Competitive mechanisms for inhibition of sulfate reduction and methane production in the zone of ferric iron reduction in sediments Appl Environ Microbiol 53:2636–2641.

72. Lovley DR, Holmes DE, Nevin KP. 2004. Dissimilatory Fe(III) and Mn(IV) reduction. Adv Microb Physiol 49:219–86.

73. Pritchett MA, Zhang JK, Metcalf WW. 2004. Development of a markerless genetic exchange method for *Methanosarcina acetivorans* C2A and its use in construction of new genetic tools for methanogenic archaea. Appl Environ Microbiol 70:1425–33.

74. Lovley DR, Phillips EJ. 1988. Novel mode of microbial energy metabolism: organic carbon oxidation coupled to dissimilatory reduction of iron or manganese. Appl Environ Microbiol 54:1472–80.

75. Mouser PJ, Holmes DE, Perpetua LA, DiDonato R, Postier B, Liu A, Lovley DR. 2009. Quantifying expression of *Geobacter spp*. oxidative stress genes in pure culture and during in *situ* uranium bioremediation. ISME J 3:454–465.

76. Holmes DE, Giloteaux L, Orellana R, Williams KH, Robbins MJ, Lovley DR. 2014. Methane production from protozoan endosymbionts following stimulation of microbial metabolism within subsurface sediments. Front Microbiol 5.

77. Lovley DR, Coates JD, BluntHarris EL, Phillips EJP, Woodward JC. 1996. Humic substances as electron acceptors for microbial respiration. Nature 382:445–448.

78. Holmes DE, Risso C, Smith JA, Lovley DR. 2012. Genome-scale analysis of anaerobic benzoate and phenol metabolism in the hyperthermophilic archaeon *Ferroglobus placidus*. ISME J 6:146–57.

79. Bolger AM, Lohse M, Usadel B. 2014. Trimmomatic: a flexible trimmer for Illumina sequence data. Bioinformatics 30:2114–20.

80. Magoc T, Salzberg SL. 2011. FLASH: fast length adjustment of short reads to improve genome assemblies. Bioinformatics 27:2957–63.

81. Kopylova E, Noe L, Touzet H. 2012. SortMeRNA: fast and accurate filtering of ribosomal RNAs in metatranscriptomic data. Bioinformatics 28:3211–7.

82. Mortazavi A, Williams BA, McCue K, Schaeffer L, Wold B. 2008. Mapping and quantifying mammalian transcriptomes by RNA-Seq. Nat Methods 5:621–8.

83. Klevebring D, Bjursell M, Emanuelsson O, Lundeberg J. 2010. In-Depth Transcriptome Analysis Reveals Novel TARs and Prevalent Antisense Transcription in Human Cell Lines. Plos One 5.

84. Robinson MD, McCarthy DJ, Smyth GK. 2010. edgeR: a Bioconductor package for differential expression analysis of digital gene expression data. Bioinformatics 26:139–40.

85. Marchler-Bauer A, Derbyshire MK, Gonzales NR, Lu S, Chitsaz F, Geer LY, Geer RC, He J, Gwadz M, Hurwitz DI, Lanczycki CJ, Lu F, Marchler GH, Song JS, Thanki N, Wang Z, Yamashita RA, Zhang D, Zheng C, Bryant SH. 2015. CDD: NCBI’s conserved domain database. Nucleic Acids Res 43:D222–6.

86. Finn RD, Coggill P, Eberhardt RY, Eddy SR, Mistry J, Mitchell AL, Potter SC, Punta M, Qureshi M, Sangrador-Vegas A, Salazar GA, Tate J, Bateman A. 2016. The Pfam protein families database: towards a more sustainable future. Nucleic Acids Res 44:D279–85.

87. Hofmann K, Stoffel W. 1993. TMBASE-A database of membrane spanning protein segments. Biol Chem Hoppe-Seyler 374:166.

88. Krogh A, Larsson B, von Heijne G, Sonnhammer EL. 2001. Predicting transmembrane protein topology with a hidden Markov model: application to complete genomes. J Mol Biol 305:567–80.

89. Tusnady GE, Simon I. 2001. The HMMTOP transmembrane topology prediction server. Bioinformatics 17:849–50.

90. Yu NY, Wagner JR, Laird MR, Melli G, Rey S, Lo R, Dao P, Sahinalp SC, Ester M, Foster LJ, Brinkman FS. 2010. PSORTb 3.0: improved protein subcellular localization prediction with refined localization subcategories and predictive capabilities for all prokaryotes. Bioinformatics 26:1608–15.

91. Petersen TN, Brunak S, von Heijne G, Nielsen H. 2011. SignalP 4.0: discriminating signal peptides from transmembrane regions. Nat Methods 8:785–6.

