## Supplementary Table S2 for "A Membrane-Bound Cytochrome Enables *Methanosarcina acetivorans* to Conserve Energy to Support Growth from Extracellular Electron Transfer"

**Supplementary Table S2**. Differential expression of genes coding for CO_2_ reduction pathway genes and methyl-coenzyme M reductase (Mcr) in *M. acetivorans* cells grown with methanol and AQDS in the presence of BES or cells grown via methanogenesis with methanol. Negative values indicate that genes were more significantly expressed in methanogenic cells. Genes were only considered differentially expressed if the fold change was ≥ 2 and the P-value and FDR (False Discovery Rate) were <0.05.

NS: no significant difference in read abundance between conditions

| Locus ID | Annotation | Gene | Fold up-regulated AQDS/BES vs methanogenesis | P-value | FDR |
| --- | --- | --- | --- | --- | --- |
| MA4546 | methyl-coenzyme M reductase, α-subunit | *mcrA* | -1.04 (NS) | 0.812 | 0.850 |
| MA4547 | methyl-coenzyme M reductase, γ-subunit | *mcrG* | -1.29 (NS | 0.131 | 0.193 |
| MA4548 | methyl coenzyme M reductase, subunit D | *mcrD* | -1.50 (NS) | 0.039 | 0.071 |
| MA4549 | methyl coenzyme M reductase, subunit C | *mcrC* | -1.32 (NS) | 0.100 | 0.154 |
| MA4550 | methyl-coenzyme M reductase, β-subunit | *mcrB* | 1.09 (NS) | 0.535 | 0.614 |
| MA0269 | THMPT S-methyltransferase, subunit H | *mtrH* | -2.51 | 1.61x10^-5^ | 0.0001 |
| MA0270 | THMPT S-methyltransferase, subunit G | *mtrG* | -4.08 | 4.16x10^-6^ | 6.60x10^-5^ |
| MA0271 | THMPT S-methyltransferase, subunit F | *mtrF* | -3.32 | 1.79x10^-6^ | 3.89x10^-5^ |
| MA0272 | THMPT S-methyltransferase, subunit A | *mtrA* | -2.90 | 6.63x10^-6^ | 9.13x10^-5^ |
| MA0273 | THMPT S-methyltransferase, subunit B | *mtrB* | -2.42 | 5.01x10^-5^ | 3.89x10^-5^ |
| MA0274 | THMPT S-methyltransferase, subunit C | *mtrC* | -2.57 | 2.32x10^-5^ | 0.0002 |
| MA0275 | THMPT S-methyltransferase, subunit D | *mtrD* | -2.49 | 3.71x10^-5^ | 0.0002 |
| MA0276 | THMPT S-methyltransferase, subunit E | *mtrE* | -2.98 | 5.53x10^-5^ | 0.0003 |
| MA3733 | methylenetetrahydromethanopterin reductase | *mer* | -2.32 | 3.67x10^-5^ | 0.0002 |
| MA4430 | methylenetetrahydromethanopterin dehydrogenase | *mtd* | -3.11 | 7.56x10^-6^ | 9.89x10^-5^ |
| MA1710 | methenyltetrahydromethanopterin cyclohydrolase | *mch* | -1.77 (NS) | 0.001 | 0.003 |
| MA0010 | formylmethanofuran-tetrahydromethanopterin formyltransferase | *ftr* | -1.48 (NS) | 0.042 | 0.074 |
| MA0304 | formylmethanofuran dehydrogenase, subunit E | *fmdE* | -1.22 (NS) | 0.139 | 0.203 |
| MA0305 | formylmethanofuran dehydrogenase, subunit F | *fmdF* | -1.05 (NS) | 0.675 | 0.737 |
| MA0306 | formylmethanofuran dehydrogenase, subunit A | *fmdA* | -1.35 (NS) | 0.050 | 0.087 |
| MA0307 | formylmethanofuran dehydrogenase, subunit C | *fmdC* | -1.47 (NS) | 0.014 | 0.031 |
| MA0308 | formylmethanofuran dehydrogenase, subunit D | *fmdD* | -1.46 (NS) | 0.016 | 0.033 |
| MA0309 | formylmethanofuran dehydrogenase, subunit B | *fmdB* | -1.12 (NS) | 0.512 | 0.591 |
| MA0975 | Coenzyme F420-reducing hydrogenase subunit alpha | *frhA* | 2.77 | 1.12x10^-5^ | 0.0001 |
| MA0976 | Coenzyme F420-reducing hydrogenase subunit delta | *frhD* | 2.67 | 5.95x10^-5^ | 0.0004 |
| MA0977 | Coenzyme F420-reducing hydrogenase subunit gamma | *frhG* | 2.35 | 4.95x10^-5^ | 0.0003 |
| MA0978 | Coenzyme F420-reducing hydrogenase subunit beta | *frhB* | 2.04 | 0.0004 | 0.001 |
| MA0687 | Heterodisulfide reductase subunit E | *hdrE* | -1.47 (NS) | 0.021 | 0.041 |
| MA0688 | Heterodisulfide reductase subunit D | *hdrD* | -1.18 (NS) | 0.303 | 0.387 |
