## Supplementary Table S3 for "A Membrane-Bound Cytochrome Enables *Methanosarcina acetivorans* to Conserve Energy to Support Growth from Extracellular Electron Transfer"

**Supplementary Table S3**. Differential expression of genes coding for proteins from the proton-translocating F_420_:methanophenazine oxidoreductase (Fpo) complex in *M. acetivorans* cells grown via methanogenesis with methanol vs cells grown with methanol as the electron donor and AQDS (in the presence of BES) as the acceptor. Genes were only considered differentially expressed if the fold change was ≥ 2 and the P-value and FDR (False Discovery Rate) were <0.05.

| Locus ID | Annotation | Gene | Fold up-regulated in methanogenesis vs AQDS/BES | log_2_ RPKM AQDS/BES | log_2_ RPKM methanogenesis |
| --- | --- | --- | --- | --- | --- |
| MA1495 | F420H2 dehydrogenase subunit A | *fpoA* | 3.76 | 5.76x10^-6^ | 8.33x10^-5^ |
| MA1496 | F420H2 dehydrogenase subunit B | *fpoB* | 3.26 | 7.15x10^-6^ | 9.50x10^-5^ |
| MA1497 | F420H2 dehydrogenase subunit C | *fpoC* | 3.56 | 3.19x10^-6^ | 5.51x10^-5^ |
| MA1498 | F420H2 dehydrogenase subunit D | *fpoD* | 3.25 | 6.94x10^-6^ | 9.34x10^-5^ |
| MA1499 | F420H2 dehydrogenase subunit H | *fpoH* | 2.61 | 0.0001 | 0.0008 |
| MA1500 | F420H2 dehydrogenase subunit I | *fpoI* | 4.28 | 3.75x10^-6^ | 6.15x10^-5^ |
| MA1501 | F420H2 dehydrogenase subunit J | *fpoJ1* | 4.43 | 1.49x10^-5^ | 0.0001 |
| MA1502 | F420H2 dehydrogenase subunit J | *fpoJ2* | 4.20 | 1.36x10^-5^ | 0.0001 |
| MA1503 | F420H2 dehydrogenase subunit K | *fpoK* | 3.71 | 4.63x10^-5^ | 0.0003 |
| MA1504 | F420H2 dehydrogenase subunit L | *fpoL* | 3.62 | 4.97x10^-6^ | 7.58x10^-5^ |
| MA1505 | F420H2 dehydrogenase subunit M | *fpoM* | 2.99 | 5.14x10^-5^ | 0.0003 |
| MA1506 | F420H2 dehydrogenase subunit N | *fpoN* | 2.77 | 0.0004 | 0.001 |
| MA1507 | F420H2 dehydrogenase subunit 0 | *fpoO* | 2.76 | 0.0002 | 0.001 |
| MA3732 | F420H2 dehydrogenase subunit F | *fpoF* | 2.77 | 8.14x10^-6^ | 0.0001 |
