## Supplementary Table S4 for "A Membrane-Bound Cytochrome Enables *Methanosarcina acetivorans* to Conserve Energy to Support Growth from Extracellular Electron Transfer"

**Supplementary Table S4:** Primers used to construct various deletion mutant strains (restriction sites are underlined). Genetically recombinant plasmids were checked for accuracy by PCR with verification primers.

| Primer name | Mutant Construct |  |
| --- | --- | --- |
| flaB1flaB2 up-f  flaB1flaB2 up-r  flaB1flaB2 down-f  flaB1flaB2 down-r | flaB1flaB2 double deletion mutant | TCTCTCGAGTTCCTTGAAGATATTAAAGGTC  TCTAAGCTTAATGAATCACCTCAATATTGTG  TCTGGATCCAGCTTGAAATCAAACCAC  TCTGCGGCCGCCACTGCAGCTATAACAC |
| flaB1flaB2 up verification  flaB2flaB2 down verification |  | ACTCTATGCTTGCAGCTGAC/TCTGCGGCCGCCACTGCAGCTATAACAC  CCGCCTGCAGTATTCGTTAC/TCTCTCGAGTTCCTTGAAGATATTAAAGGTC |
| Verification of replacement of flaB1flaB2 with pac gene |  | AGAGACCCTATCTTACCTGC/TCTGCGGCCGCCACTGCAGCTATAACAC |
| MA0658 up-f  MA0658 up-r  MA0658 down-f  MA0658 down-r | MA0658 deletion mutant | TCTGAGCTCGATAAAAGCTTGTCATGC  ATCTCTAGACAGCCTCCGCAGACCGAATC  TCTGAATTCGGAATGAAGCCGTGATTGAC  TCTCTCGAGTCCCGAACTTGACAATTAC |
| MA0658 deletion verification |  | CTAGTCGTTCATGCCAGTTC/AGGACGTTAGCAGTGTAGAC |
| Verification of pac gene replacement |  | CTAGTCGTTCATGCCAGTTC/AGTATATTACGAATAGGGCG |
| MA3739 up-f  MA3739 up-r  MA3739 down-f  MA3739 down-r | MA3739 deletion mutant | TCTGTCGACTATCGGAGTAAGGAGACTGG  ATCTCTAGACTCCCAGGTGCACGTTTGC  TTCGAGCTCCACAGGAACGCTTATTGC  TCTGCGGCCGCTACCAAAATAACACCCTCC |
| MA3739 deletion verification  primers |  | TCTGTCGACTATCGGAGTAAGGAGACTGG/TGTCCGCATTCGAATGCATC |
| Verification of pac gene replacement |  | TCTGTCGACTATCGGAGTAAGGAGACTGG/AGTATATTACGAATAGGGCG |
| MA2908 up-f  MA2908 up-r  MA2908 down-f  MA2908 down-r | MA2908 deletion mutant | ATGCTCGAGTCTGGTGGTAACAGGACTA  ACGAAGCTTCTACCATCTACTTCTAAACTAC  ATGGGATCCCAATTGTTGCCTTCCTGCT  ATGCGGCCGCGAAGAAAGTACCTTACTGC |
| MA2908 deletion verification primers |  | ACCCTCTGGAAATGCACAAC/TAAACTGCAATCGCATCAGC |
| Verification of pac gene replacement |  | TCTGGTGGTAACAGGACTA/AGTATATTACGAATAGGGCG |
| MA0167 up-f  MA0167 up-r  MA0167 down-f  MA0167 down-r | MA0167 deletion mutant | ATGCTCGAGGAAAAGTTCAGTTCCATTC  ATGAAGCTTCAGGAATGGATTGCTTTTGC  ATGGGATCCGTCGTTGAATACCTGAAAAC  ATGCGGCCGCAGTTTTTAAGCTTCCGA |
| MA0167 deletion verification primers |  | CATTGGCTGTACGTTCATGG/CAACGACGTCATTCACATCC |
| Verification of pac gene replacement |  | GAAAAGTTCAGTTCCATTC/AGTATATTACGAATAGGGCG |
| MA2925 up-f  MA2925 up-r  MA2925 down-f  MA2925 down-r | MA2925 deletion mutant | ATGCTCGAGGACTCCCTGATTAACCTGGA  GTGAAGCTTTATTCTGGTTTATCCCGGAA  ATGGGATCCACTCTCTCGGACGGCTACTA  ATGCGGCCGCCACGATAAACAACAACAT |
| MA2925 deletion verification primers |  | CCGATGAGAATAAGGGCAAA/CGTCCGAGAGAGTTTTCAGG |
| Verification of pac gene replacement |  | GACTCCCTGATTAACCTGGA/AGTATATTACGAATAGGGCG |
