## Supplementary Figure S1 for "A Membrane-Bound Cytochrome Enables *Methanosarcina acetivorans* to Conserve Energy to Support Growth from Extracellular Electron Transfer"

### Slide 1
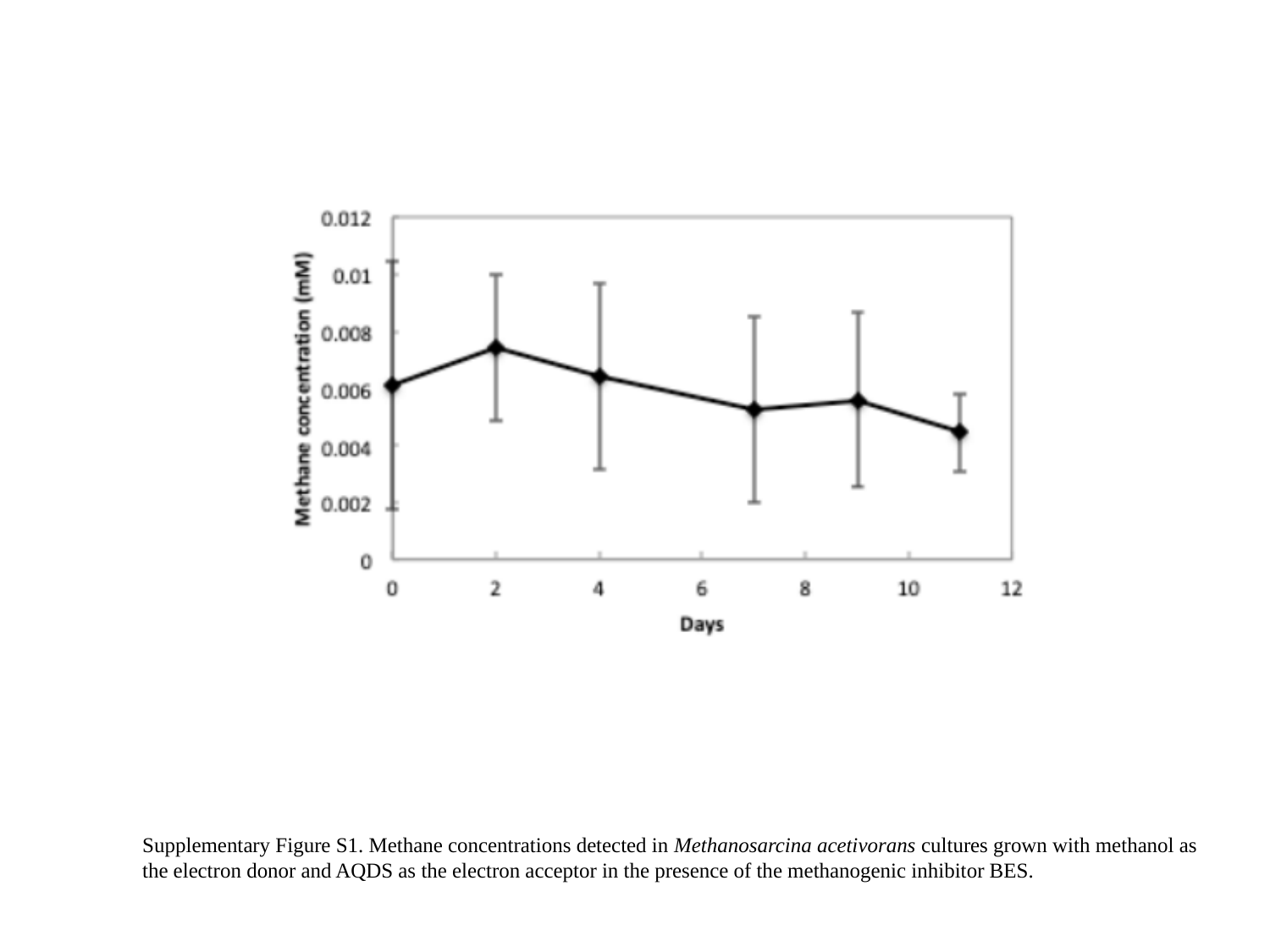

Supplementary Figure S1. Methane concentrations detected in Methanosarcina acetivorans cultures grown with methanol as the electron donor and AQDS as the electron acceptor in the presence of the methanogenic inhibitor BES.
