## Supplementary Figure S2 for "A Membrane-Bound Cytochrome Enables *Methanosarcina acetivorans* to Conserve Energy to Support Growth from Extracellular Electron Transfer"

### Slide 1
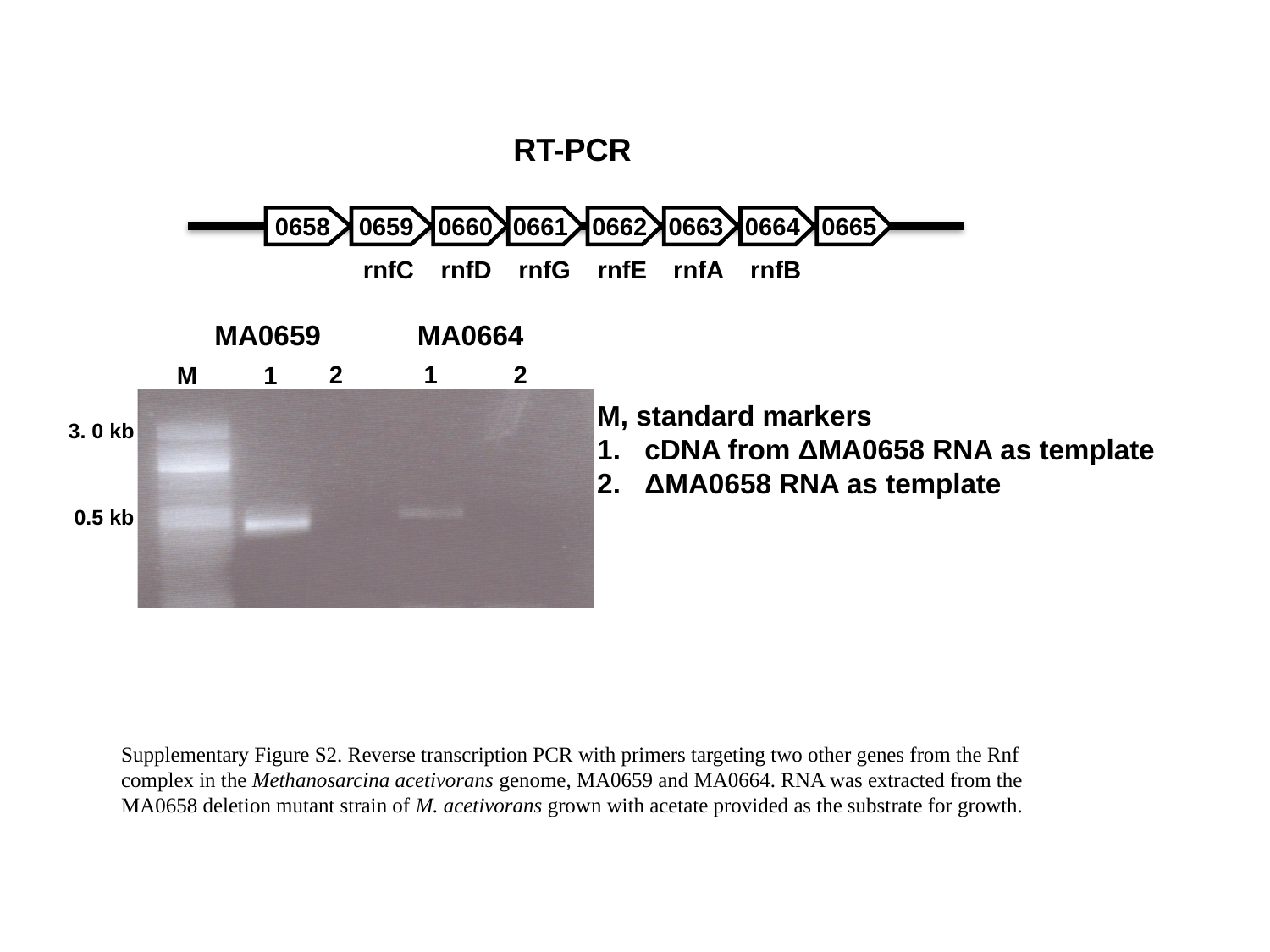

RT-PCR
0658
0659
0660
0661
0662
0663
0664
0665
rnfC
rnfD
rnfG
rnfE
rnfA
rnfB
MA0659
MA0664
2
1
2
M
1
M, standard markers
cDNA from ΔMA0658 RNA as template
ΔMA0658 RNA as template
3. 0 kb
0.5 kb
Supplementary Figure S2. Reverse transcription PCR with primers targeting two other genes from the Rnf complex in the Methanosarcina acetivorans genome, MA0659 and MA0664. RNA was extracted from the MA0658 deletion mutant strain of M. acetivorans grown with acetate provided as the substrate for growth.
